# Gosha: a database of organisms with defined optimal growth temperatures

**DOI:** 10.1101/2021.12.21.473645

**Authors:** Karla Helena-Bueno, Charlotte R. Brown, Sergey Melnikov

## Abstract

Currently, we are witnessing an explosive accumulation of genomic sequences for organisms across all branches of life. However, typically the genomic data lack the information about optimal growth conditions of corresponding organisms. As a result, it becomes challenging to use the genomic data for studying the adaptations of organisms and biological molecules to diverse environments. To address this problem, we have created a database Gosha, available at http://melnikovlab.com/gshc. This database brings together information about the genomic sequences and optimal growth temperatures for 25,324 species, including ∼89% of the bacterial species with known genome sequences. Using this database, one can annotate genomic sequences from thousands of species and correlate variations in genes and genomes with optimal growth temperatures. The database interface allows users to retrieve optimal growth temperatures for bacteria, eukaryotes and archaea, providing a tool to explore how organisms, genomes, and individual proteins and nucleic acids adapt to certain temperatures. We hope that this database will contribute to medicine and biotechnology by helping to create a better understanding of molecular adaptations to heat and cold, leading to new ways to preserve biological samples, engineer useful enzymes, and develop biological materials and organisms with the desired tolerance to heat and cold.

**GRAPHICAL ABSTRACT:** Gosha (available at www.melnikovlab.com/gshc) is a database that collects information about the optimal growth temperatures of living species. This database aims to facilitate studies of molecular adaptation to specific temperatures.

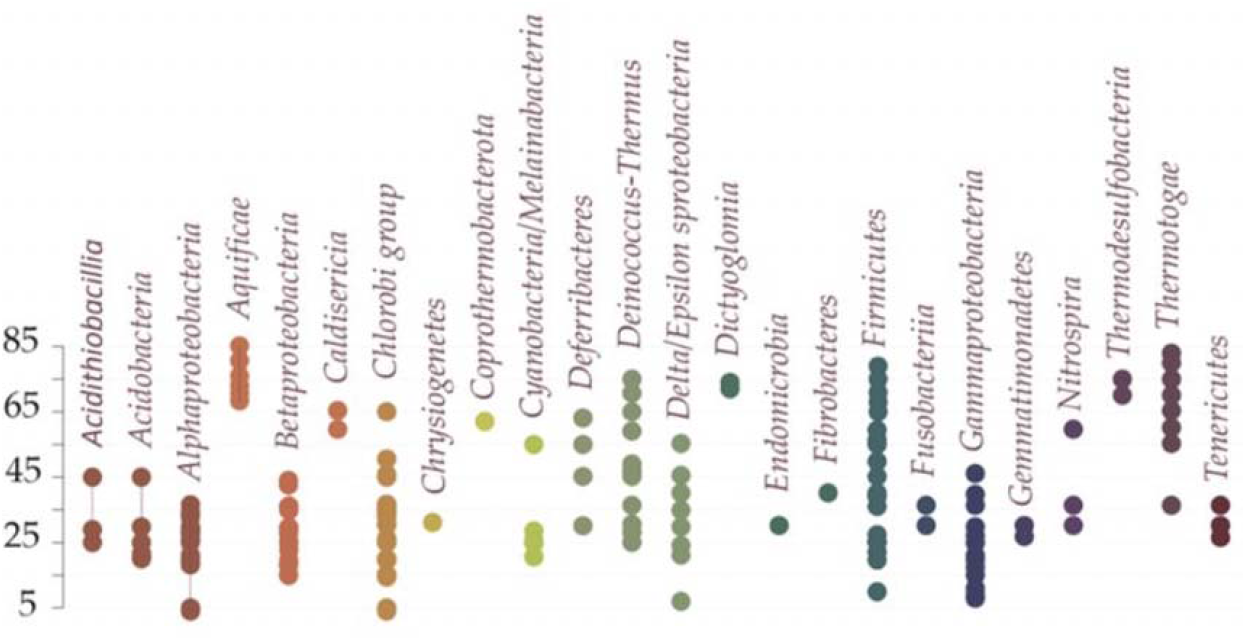

## INTRODUCTION

Despite significant research efforts to understand how biological molecules adapt to temperature change (1-36), we are still not able to accurately answer two fundamental questions: What are the most common strategies by which cellular proteins adapt to environmental conditions, such as heat and cold? And can we find a simple and robust approach to alter the thermal tolerance of natural proteins by introducing a minimal number of mutations to the protein sequence?

One challenge in answering these questions stems from the lack of a resource that stores easy-to-use information about the optimal growth conditions of living organisms, together with their genomic data. For example, currently, there are more than 14,400 genome sequences from representative bacterial species publicly available. In principle, we could use these sequences to study thousands of variants of a given protein, observing how its sequence and structure undergo changes upon transition from cold-adapted bacteria (10-16,37-48) to heat-adapted bacteria (26-32,49-69).

In practice, however, it is not immediately possible to organise thousands of organisms (along with their genomic sequences) by their optimal growth temperatures, because the corresponding genomic sequences deposited in public repositories (such as NCBI Genomes) lack information about these organisms’ optimal growth conditions. To overcome this challenge, several teams have dedicated their efforts to annotating the existing genomic sequences with experimentally defined growth conditions. Initially, lists of optimal growth conditions were compiled by conducting manual searches through the Bergey’s manual of systematic bacteriology (70,71). Later, advancements in automation made it feasible to retrieve data from thousands of species, enabling the creation of a database encompassing 8,645 prokaryotic strains with known genome sequences (72,73). However, despite having access to genome sequences for tens of thousands of representative bacteria, eukaryotes, and archaea, our current capability to efficiently categorize their genomic sequences) by optimal growth conditions remains limited, which hinders our progress in understanding the molecular and organismal adaptation to temperature.

Here, we take one step forward in developing such a resource for scientists and engineers interested in exploring and exploiting molecular adaptations to heat and cold. Using the NCBI database of sequenced genomes as a scaffold, we have created a database Gosha (<u>G</u>enomic sequences and <u>O</u>ptimal growth temperatures <u>S</u>how <u>H</u>eat-and-cold <u>A</u>daptations) in which species with known genome sequences are annotated with information about these species’ optimal growth temperature. This database describes the optimal growth temperature of more than 25,000 microorganisms, which significantly increases the volume of the database and includes 89% of the representative bacteria whose genome sequences are deposited in the NCBI database.

With regards to bacterial species, this new resource makes it possible to retrieve up to 12,354 sequences of a given bacterial protein of interest, sort these sequences by the optimal growth temperature of its corresponding species, and explore how each residue in this protein varies in sequence and conservation upon transition from cold-adapted to heat-adapted organisms. Thus, we provide a tool for large-scale studies of the molecular and organismal adaptations to a specific temperature.

## DATABASE FEATURES

### Downloadable lists of species with known genome sequence and together with their optimal growth temperature

Currently, the Gosha database contains information about the optimal growth temperature for 25,324 species. This information is continuously retrieved by web-scraping 23 public repositories of microorganisms (**Table 1**) and then added to the list of organisms deposited in the NCBI repository of organisms with sequenced genomes. The Gosha site contains the optimal growth temperatures for an organism; however, it does not yet include information about other growth conditions, such as oxygen requirement, pH, pressure, and salt concentration.

**Table 1.**
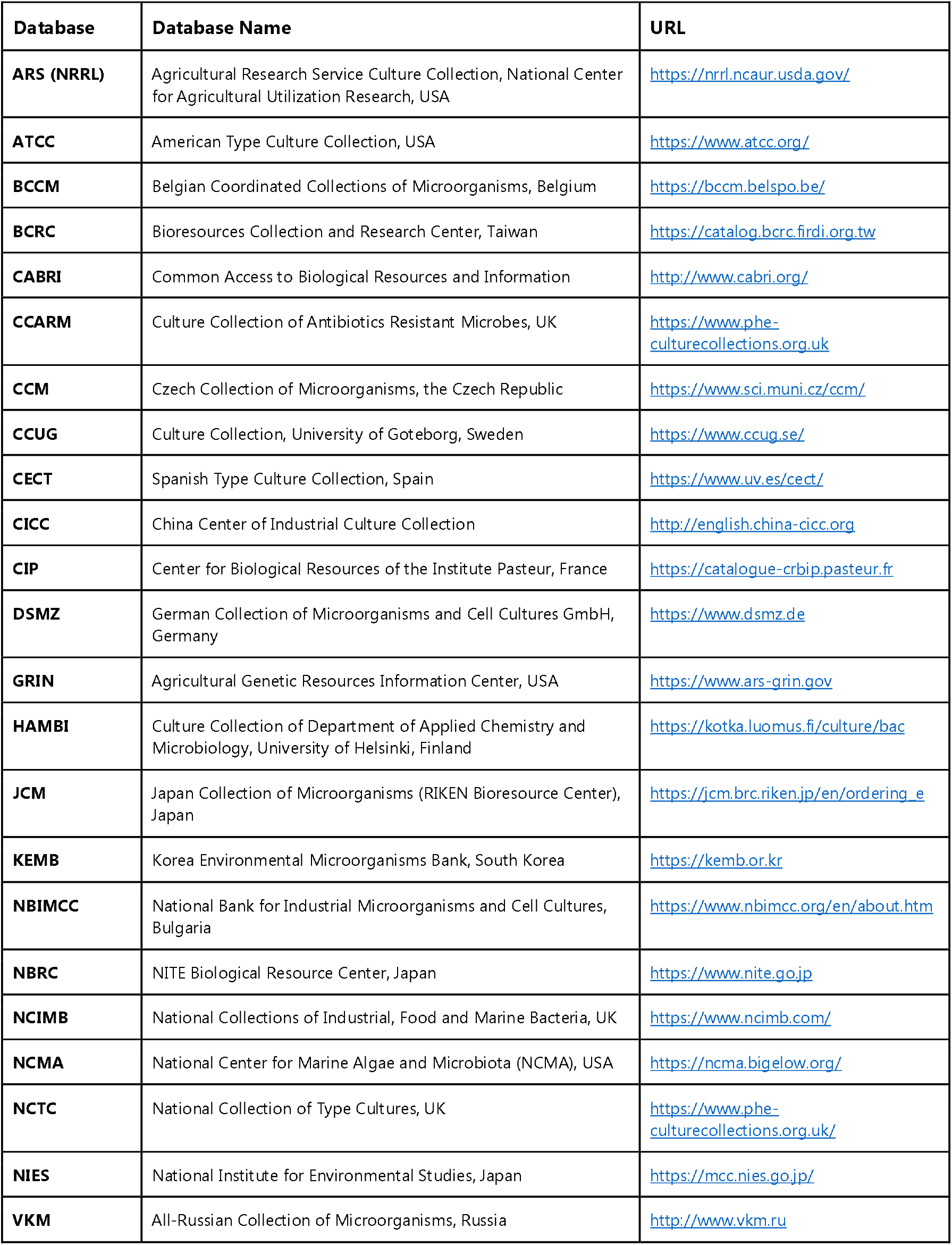
Websites used for web scraping to collect information about the optimal growth temperatures of microbial organisms.

Lists of organisms with experimentally defined optimal growth temperatures and a reference to the corresponding genome data can be downloaded from the database. This includes the optimal growth temperature values for 12,265 representative bacteria, 414 representative archaea, and 973 representative eukaryotes. The datasets are updated monthly and are available in .csv format, enabling the species to be sorted by an organism’s name, phylogenetic group, genome size, genomic GC-content, number of protein coding genes, and optimal growth temperature. As shown in the example provided for bacterial species (**Figure 1**), organisms contained in the datasets include thermophiles and psychrophiles from all major lineages of species, providing an opportunity to study molecular adaptation to heat and cold at the scale of the entire tree of life.

**Figure 1.**
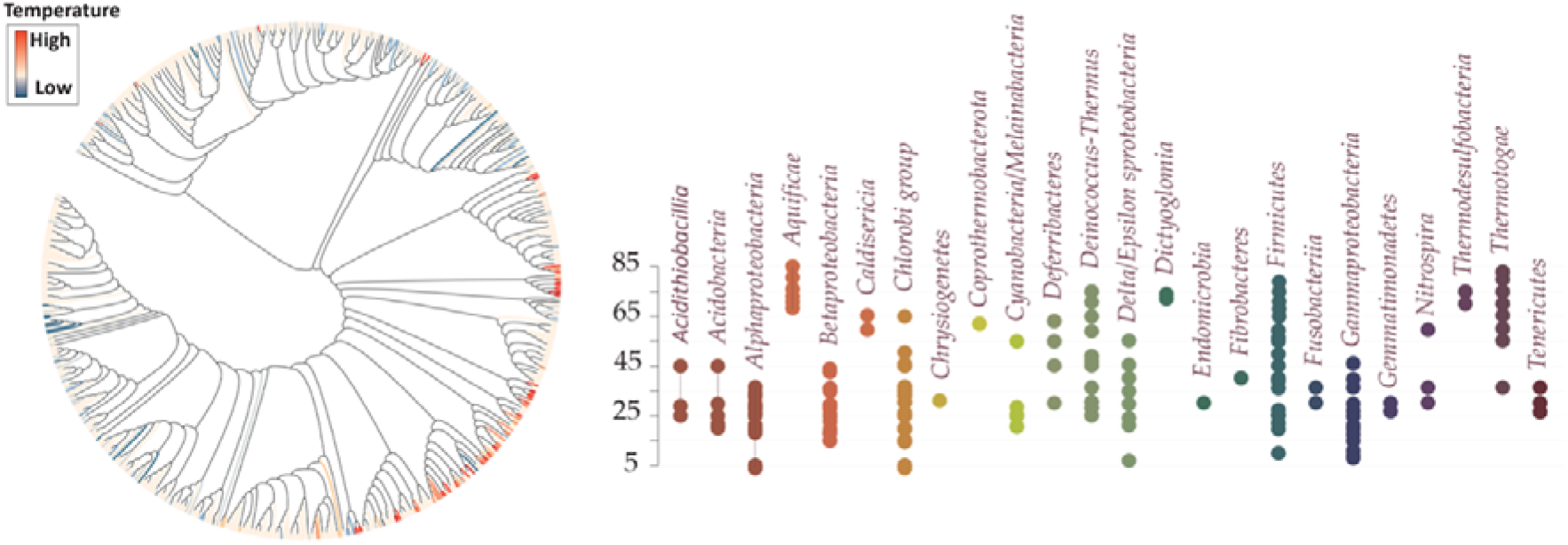
Using the database to colour bacterial species by their optimal growth temperatures. The bacterial tree of life shows representative members of bacterial genera, with each organism coloured by the optimal growth temperature ranging from 4°C (“low”) to 85°C (“high”). The dot-plot shows lineages of bacterial species in the database and their corresponding optimal growth temperature.

### Optimal growth-temperature checker

In addition to the downloadable data, the database user interface allows searching for the optimal growth temperature for a given species. The user can enter a species name in a search window and retrieve the optimal growth temperature for the species of interest if it is present in the database.

### Error and request tracking

To document the rapidly expanding data, most entries in the database are automated, without inspection against published literature. We acknowledge that the optimal growth temperature values for some organisms may contain discrepancies or inaccuracies and therefore encourage users to provide feedback on any unaddressed discrepancies by submitting reports through our feedback tracking system. The submission form allows users to request changes in the database and monitor the progress of each inquiry.

## APPLICATIONS

How will this database help researchers to explore molecular mechanisms of thermal adaptation? Below, we propose some example applications:

First, we can now sort genomes or homologous gene sequences by their optimal growth temperature (and not just by phylogenetic origin), making it possible to explore universal strategies of adaptations to heat and cold, as opposed to idiosyncratic adaptations within each lineage of species (**Figure 1**). Such studies would have far-reaching implications in biotechnology as they can simplify the rational design of biological molecules and organisms with a desired thermal stability (32,61,68,74-79).

Second, this database may simplify studies of molecular adaptations to temperature within any given range of temperatures. This is important, because to date most studies have focused on extremophiles (49,50), leaving mostly unexplored how mesophilic organisms adapt to relatively subtle changes in the environment (e.g. temperature increases of a few degrees Celsius as a consequence of climate change (12,20,21,80).

Third, we can monitor how the identity and conservation of each residue in a protein of interest gradually changes across a range of optimal growth temperatures from 2°C to more than 103°C, simplifying studies of structural constraints (21,46,63,79,81-84), and finding new ways to engineer useful proteins with a desired optimal thermal tolerance (24,76-79,84,85) (**Figure 2**).

**Figure 2.**
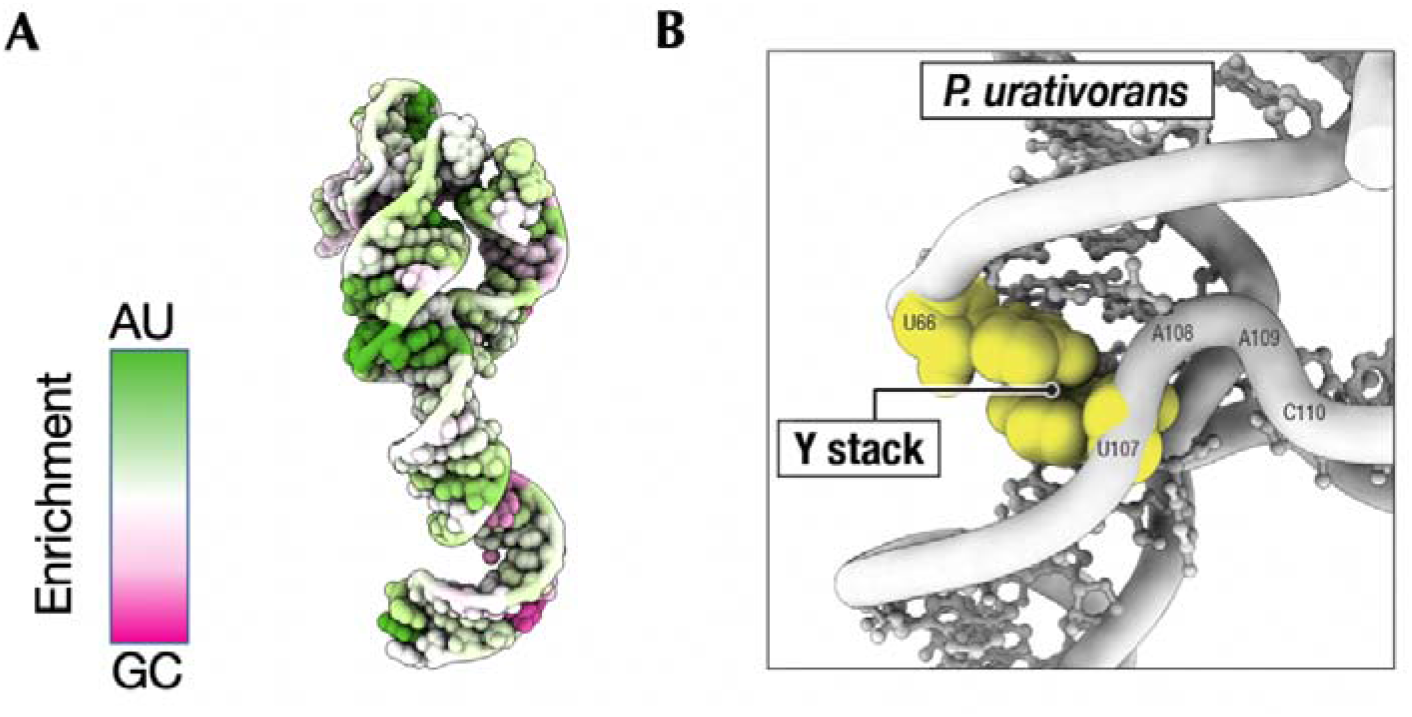
Using the database to colour molecules by thermal adaptation and identify psychrophile-specific variations that may help RNA molecules adapt to cold environments. **a**. The molecular structure of 5S rRNA from the psychrophilic bacterium *Psychrobacter urativorans* is color-coded based on the GC-bias of its residues in psychrophilic species. Residues coloured in green indicate a high preference for mutations from G or C to A or U in psychrophilic bacteria compared to thermophilic bacteria. Residues coloured in magenta highlight residues that exhibit the highest rate of mutation from A or U to G or C in psychrophilic bacteria compared to thermophilic bacteria. **b**. The atomic structure zooming in on the 5S rRNA segment with the highest AU-bias in psychrophilic bacteria reveals the presence of an unusual structural motif in psychrophilic rRNA. This motif is stabilized by the stacking of two uracil bases (U66 and U107), which is a weak interaction that differs from the typical tertiary structures of 5S rRNA in mesophilic and thermophilic bacteria, where this stacking usually involves at least one purine base to form a stronger contact.

Fourth, this database can help identify model organisms to observe “extremophiles in the making”. Currently, the database contains organisms from the same genus that have almost identical sequences for most of their cellular proteins but exhibit dramatically different optimal growth temperatures. For example, the genus *Clostridium* includes species with an optimal growth temperature ranging from just 5°C (*Clostridium frigoris*) (86), to 55°C (*Clostridium thermobutyricum*) (87). Comparing species in these genera can provide us with a rare opportunity to observe the natural transformation of mesophiles into extremophiles and gain an understanding of how organisms evolve the ability to tolerate heat and cold through minimal changes in their genomes (8,13,32,55,57,69,88). These studies are important, as they may help to simplify the design of economically useful microorganisms with the desired thermal tolerance.

## FUTURE DIRECTIONS

We are currently working to expand the database by including additional environmental parameters, such as optimal salt concentration, pH, pressure, and oxygen requirement. We are also testing a new data scraping algorithm to retrieve data not only from repositories of commercially available microorganisms but also from original research papers, to maximise the completeness of our datasets. Finally, we are testing scripts to allow the mapping of temperature-dependent variations in protein sequences to corresponding three-dimensional protein structures that are available in the Protein Data Bank. We hope that in the future our database or similar annotations will be integrated into centralised repositories of genome sequences, such as NCBI, making it possible to explore organismal adaptations to changing environments using the rapidly expanding collection of genomic sequences for both living and extinct species on our planet.

## DATA AVAILABILITY

The data described in this study were deposited in Figshare with the following permanent and citable DOI: https://doi.org/10.6084/m9.figshare.23257940.v1

## ACKNOWLEDGEMENTS

We acknowledge Yuri Polikanov for his encouragement that enabled this work to happen. We are also grateful to the members of the Melnikov lab for their contributions, as well as the reviewers of this manuscript, for their critical feedback and valuable suggestions to improve our work and service.

